# Toxicity of extracellular alpha-synuclein is independent of intracellular alpha-synuclein

**DOI:** 10.1101/2022.03.31.486401

**Authors:** Yanina Dening, Theresa Straßl, Viktoria Ruf, Petra Dirscherl, Alexandra Chovsepian, Alicia Stievenard, Amit Khairnar, Felix Schmidt, Florian Giesert, Jochen Herms, Johannes Levin, Marianne Dieterich, Peter Falkai, Daniela Vogt Weisenhorn, Wolfgang Wurst, Armin Giese, Francisco Pan-Montojo

## Abstract

Parkinson′s disease (PD) pathology progresses throughout the nervous system affecting numerous neuronal structures. It has been postulated that the progression of the pathology is based on a prion-like disease mechanism partly due to the seeding effect of endocytosed alpha-synuclein (ASYN) on the endogenous ASYN. The appearance of the pathology in dopaminergic neurons leads to neuronal cell death and motor symptoms. However, the effect on other neuronal structures is more inconsistent, leading to a higher variability in the prevalence of non-motor symptoms. Thus, the sensitivity to the pathology seems to vary among neuronal subtypes. Here, we analyzed the role of endogenous ASYN in the progression of PD-like pathology and the effect of monomeric and oligomeric ASYN as well as paraquat and rotenone on primary enteric, dopaminergic and cortical neurons from wild-type mice. Our results showed that pathology progression did not occur in the absence of endogenous ASYN and that dopaminergic neurons were more sensitive to ASYN and rotenone when compared to all other neuronal subtypes. Remarkably, the toxic effect of ASYN was independent of the presence of endogenous ASYN and directly related to the disturbance of the mitochondrial membrane potential. Thus, we suggest that the interaction between ASYN and mitochondria plays an important role in the toxicity of trans-synaptically transported ASYN and in the progression of PD pathology. These results question the prion-disease hypothesis and propose that endocytosed ASYN impairs the host′s mitochondrial function thereby also contributing to PD-pathology progression.

## Introduction

Alpha-synuclein (ASYN) is the main component of Lewy bodies (LB) and Lewy neurites (LN). LB and LN are not exclusive of PD [1] and mainly consist in intraneuronal and intra-glial aggregates. In idiopathic PD most of the patients that show PD related inclusions in CNS sites already present LB and LN in the ENS and the sympathetic ganglia [2]. Based on autopsies performed on PD patients and healthy individuals, Braak and colleagues developed a pathological staging of the disease [3]. According to this staging, PD lesions follow a spatio-temporal pattern that starts in the olfactory bulb (OB) and the ENS progressing into the CNS through synaptically connected structures (e.g. the enteric nervous system (ENS), the sympathetic cervical ganglia, the intermediolateral nucleus of the spinal cord (IML), the motor nucleus of the vagus (DMV) or the amygdala) [4, 5]. This pathological staging of the disease seems to correlate well with the appearance of early non-motor symptoms in PD patients. These include hyposmia, gastrointestinal alterations, autonomic dysfunction and the experience of pain [6]. Interestingly, the prevalence of non-motor symptoms is very low compared to motor symptoms with big variations between symptoms (e.g. prevalence of constipation 46.5% range: 27.5–71.7%; swallowing 25.4% range: 16.1-30.3%; REM sleep behaviour disorder 34.2% range: 29.6–38.7%)[7]. The reason for this high variability between neuronal subpopulations is still unknown. It has been postulated that dopaminergic neurons might have a higher susceptibility due to a higher metabolic demand and dopamine synthesis and metabolism leading to higher oxidative damage to mitochondria (reviewed in [8]).

The molecular mechanisms underlying the progression of PD pathology are also still unclear. Pathological studies on PD patients that had received cellular transplants show that transplanted cells display the pathology in the form of ASYN inclusions some years after transplantation. Using cell cultures and in vivo experiments with mice, it has been shown that ASYN can be transmitted between neurons and interact with the host′s ASYN and mitochondria. However, the role of these interactions in the progression of PD pathology is still unclear. It has been shown that aggregated ASYN can interact with native endogenous ASYN and it has been postulated that it acts as a seeding agent inducing the aggregation of endogenous ASYN. On the other hand, it has been shown that ASYN oligomers can inhibit the activity of mitochondrial respiratory complexes such as Complex I [9].

We have shown that orally administered rotenone induces ASYN accumulation in most of the regions described in Braak′s staging [10-12]. The appearance of these alterations followed a spatio-temporal pattern similar to the one predicted in Braak′s staging and the resection of the nerves connecting the gut to the substantia nigra pars compacta (SNc, i.e. the vagus and sympathetic nerves) prevented the progression of the pathology to the DMV and IML. This animal model also shows at least one non-motor symptom associated with PD (i.e. constipation), which was related to the loss of sympathetic and parasympathetic innervation of the intestinal tract [13, 14].

In the current study, we tested the seeding hypothesis by investigating the effect of orally administered rotenone on ASYN KO mice and dissect the role of ASYN-ASYN and ASYN-mitochondria interactions by comparing the effect of extracellular ASYN monomers and oligomers on dopaminergic neurons in primary mesencephalic neuronal cultures from wild-type, ASYN knock-out, TH-GFP and TH-GFP/ASYN knock-out mice. Finally, in order to confirm neuronal subtype specific sensitivity to external *noxae*, we compared the sensitivity of several neuronal subtypes to paraquat, rotenone and ASYN oligomers.

## Materials and methods

### Animal experiments

Male and female mice of the C57BL/6 (WT) or B6;129×1-*Snca*^*tm1Rosl*^/J (ASYN KO) strains were purchased from Jackson Laboratories (USA) over Charles River (Europe) and kept in the experimental animal facility until they reached the age to start experiments. The mice were group-housed (4-5 littermates per cage) in standard rodent polycarbonate independently ventilated cages. At the age of one year, the animals were randomly assigned to the rotenone or vehicle groups, using simple randomization [15] and the numbers were assigned using tail marks. For the experiments on ASYN KO mice, 7 mice per group (Vehicle 2 months; Rotenone 2 months; Vehicle 4 months; Rotenone 4 months) were used. The groups were treated in parallel, and rotenone exposure took place in a rotating order in the morning. The WT mice were treated following the same experimental design. Details on the experimental conditions and the results obtained can be found here [16]. Exclusion criteria included sickness (unrelated to rotenone exposure), persistent weight loss or death before the end of experiment and no mice needed to be excluded from the study.

Environmental conditions during the whole study were constant: relative humidity 50-60 %, room temperature 23 ºC ± 1 ºC, regular 12-hour light-dark cycle (6 a.m. to 6 p.m. darkness). Food and water were available *ad libitum*. All procedures were performed in accordance with EU Directive no. 2010/63/EU and approved by the Bavarian (Act. No.: 55.2-1-54-2532-5-2016) and Czech Governmental Animal Care Committee, Act No. 246/1992).

### Evaluation of motor function with Rotarod

A TSE accelerating RotaRod 5589 (TSE Systems GmbH, Germany) was used to measure motor coordination and endurance as previously described [17]. The rotarod test was always performed in morning hours, between 08:00 and 11:00 AM. Mice were placed on the cylinder and allowed to walk for 10 sec with a constant speed of 4 rpm. Gradual acceleration from 4 to 40 rpm over 5min followed and the latency to fall (sec) from the cylinder was recorded. The minimum threshold for recording rotarod activity was 10 seconds. The average performance, measured as the mean latency between the three trials, was used for analysis. As the TSE software gives automated measurements for each experiment, no blinding was required.

### Primary neuronal cultures

Primary cortical, mesencephalic and enteric neuronal cell cultures were prepared as previously described with slight modifications [11]. Brains from 6-8 E14.5 or E15.5 embryos were obtained from pregnant C57JBL6 and B6;129×1-*Snca*^*tm1Rosl*^/J mice or from B6.B6D2-Tg(Th-EGFP)21-31Koba (kindly provided by the RIKEN institute) [18] and B6.B6D2-Tg(Th-EGFP)21-31Koba/129×1-*Snca*^*tm1Rosl*^/J mice expressing TH-GFP (see Supplementary Figure 1 for genotyping) after cervical dislocation. The midbrain or the cortex (previously removing the hippocampus) were dissected under the microscope. After the dissection, the midbrain or cortical tissue from the 6-8 embryos were pooled together before digestion with Trypsin-EDTA 0.12% (Life Technologies, USA) for 7 min. The trypsin reaction was then stopped by adding basic medium (BS) containing Neurobasal A medium (Gibco), 1 mg/mL Pen/Strep, 10% FCS and 200 mM L-Glutamine and cells were mechanically dissociated using a fire-polished Pasteur pipette. Medium was fully replaced after 5 min, centrifugation at 1200 rpm, aspiring the supernatant and adding 8 mL of fresh BS to the pellet. Concentration of cells in the medium was estimated using a Neubauer chamber and a 100 μL of medium containing 1×10^6^ cells /mL plated per well in a 96-well plate (Greiner Sensoplate, Germany) coated with Poly-D-Lysine (Sigma-Aldrich, cat. no P7280) (mesencephalic) or Poly-L-Lysine (Sigma-Aldrich, cat. no P6282) (cortical). Then a 20 μL of medium was removed from the well and 24 h later 1/3 of the media was replaced with fresh BM. On DIV3 half of the medium was replaced with B27 one, containing Neurobasal A medium, 1mg/mL Pen/Strep, 200 mM L-Glutamine and B-27 supplement and on DIV5 all medium was replaced by B27 medium. Treatment was administered on DIV7 and DIV 9 cell and cells were fixed on DIV10.

Enteric and sympathetic neurons were obtained as previously described [11, 13] and enteric neurons were treated with the same protocol as cortical and dopaminergic neurons.

For the enteric neuronal culture, small intestines from 6-8 newborn mice were pooled together before dissociation with trypsin and plating took place. Briefly, the muscle layer of the gut was extracted using a forceps n.4 (FST, Germany, EU) and placed in Hank’s buffer solution (HBSS) (Invitrogen, Germany, EU). The tissue was then incubated in Collagenase I (Worthington, USA) for 45 minutes. The remaining three-dimensional enteric plexuses from the muscle layer were removed using a 200 μl pipette (Eppendorf, Germany, EU) and placed in HBSS. Enteric plexuses were incubated in trypsin-EDTA (Invitrogen, Germany, EU) for 15 minutes. The trypsin reaction was then stopped with the addition of fetal calf serum (FCS) containing media. The neurons were subsequently mechanically disaggregated and centrifuged at 1200 r.p.m. for 5 min. The supernatant was replaced by cell culture medium containing Neurobasal-A (Invitrogen, Germany, EU), 10% FCS (Sigma-Aldrich, Germany, EU), 2 mM L-Glutamine (Biochrom, Germany, EU), 1% N-2 supplement (Invitrogen, Germany, EU) 200 Units/ml penicillin (Invitrogen, Germany, EU) and 200 g/ml streptomycin (Invitrogen, Germany, EU) and cells were plated on collagen (Rat tail collagen, type I, BD Bioscience, USA) coated coverslips at the desired concentration.

For the sympathetic neuronal culture, sympathetic ganglia from 6-8 newborn mice were pooled together. Briefly, sympathetic ganglia were extracted using a forceps n.4 (FST, Germany, EU) and placed in Hank’s buffer solution (HBSS) (Invitrogen, Germany, EU). The tissue was then incubated in Collagenase I (Worthington, USA) for 20 minutes. After collagenase incubation, sympathetic ganglia were incubated in trypsin-EDTA (Invitrogen, Germany, EU) for 10 minutes. The trypsin reaction was stopped with the addition of fetal calf serum (FCS) containing media. The neurons were subsequently mechanically disaggregated and centrifuged at 1200 r.p.m. for 5 min. The supernatant was replaced by cell culture medium containing Neurobasal-A (Invitrogen, Germany, EU), 10% FCS (Sigma-Aldrich, Germany, EU), 2 mM L-Glutamine (Biochrom, Germany, EU), 1% N-2 supplement (Invitrogen, Germany, EU) 200 Units/ml penicillin (Invitrogen, Germany, EU), 100ng/ml 2.5S Nerve growth factor (NGF; BD Biosciences, cat. no. 354005) and 100 μg/ml penicillin/streptomycin (Invitrogen, Germany, EU) and cells were plated on a lateral compartment of the Campenot chambers, as previously described [19], at the desired concentration. ASYN oligomers were only added to the neurite compartment.

For all *in vitro* experiments, contaminated wells were excluded from the study

### Induction of ASYN fibrillation

ASYN oligomers for the experiments with dopaminergic neurons were prepared as previously described [20]. Briefly, with an amino reactive fluorescent dye (Alexa Fluor-647 or Alexa Fluor-488) fluorescently labelled recombinant human-ASYN and unlabelled human-ASYN [21] were mixed at a ratio of 1:5. Oligomer formation was induced at a concentration of 50 μM ASYN in the presence of 100 μM Al^3+^ under constant shaking (1400 rpm) at 37 °C, over 30 minutes. Oligomer formation was confirmed by single molecule fluorescence spectroscopy [22] via the fluorescently labelled ASYN molecules. Oligomers were aliquoted and stored until use at -80 °C.

### Drugs and treatments

#### Oral rotenone treatment

Rotenone was obtained from Sigma-Aldrich (Merck, cat. no 45656) and dissolved in chloroform (50 mg in 1 ml). This solution was kept at -20°C and used to prepare the final solutions for administration during the study. The final rotenone solution was prepared fresh every 3^rd^ day before the administration as follows. 100 microliters of the stored rotenone solution (containing 5 mg of rotenone) was added to 8 ml of 2% carboxymethylcellulose solution to get 0.625 mg/ml rotenone solution. An oral gavage was used to administer 0.01 ml/g animal weight of rotenone solution. Carboxymethylcellulose solution 2% with 1.25% chloroform was used as a vehicle in control animals. This animal model was generated in parallel to 1-year old WT mice (protocol and results published in [16]) by administration via intragastric gavage 5 days per week for 2 or 4 months (see Supplementary Figure 2). The oral administration was always performed during the morning hours between 09:00 and 11:00 AM.

### Rotenone, paraquat and ASYN treatment

Treatment with rotenone, paraquat and ASYN oligomers or monomers took place on DIV7 and DIV9. Rotenone was previously dissolved in 100% ethanol to a concentration of 10mM and stored as stock solution at -20°C, paraquat (Merck, cat. no 36541) was dissolved in water to a concentration of 1M and stored at -20°C as stock solution. ASYN oligomers and monomers were prepared as previously described.

On DIV7 rotenone, paraquat or ASYN were diluted to a final concentration of 20 nM, 25 μM or 12.5 μM and 2 or 0.2 μM respectively and half of the medium in each well was replaced by the freshly prepared medium containing the different substances. On DIV9 rotenone, paraquat or ASYN were diluted to a final concentration of 50 nM or 10 nM, 12.5 μM or 6.25 μM and 1 or 0.1 μM respectively and the medium in each well was replaced by the freshly prepared medium containing the different substances.

### Live-imaging of TH-GFP dopaminergic neurons with or without 647-tagged human ASYN

Mesencephalic neurons containing TH-GFP WT and TH-GFP ASYN KO dopaminergic neurons were stained with Tetramethylrhodamine, ethyl ester (TMRE), which is a cell-permeant, cationic, red-orange fluorescent *dye* that is readily sequestered by active mitochondria in living cells. Cells were then imaged with the help of a confocal Leica microscope (Leica TCS SP5, Leica-Microsystems, Germany) and 63X glycine objective inside a live cell-imaging chamber (37°C and 5% CO_2_) adapted to the microscope. Using the following lasers for excitation (458, 561 and 633 nm), both excitation and emission wavelengths were adjusted to obtain the best signal and avoid bleed-through between channels. In the case of sympathetic neurons in the Campenot chamber, we used Mitotracker (MitoTracker® Deep Red FM (abs/em ∼644/665 nm) as a second marker for mitochondrial membrane potential because ASYN monomers and oligomers were tagged with Alexa-Fluor-488 and because it allows fixation without losing signal intensity.

### Image analysis of TMRE and TH-GFP signals

Images obtained with the confocal microscope were analyzed using FIJI software. TH-GFP-WT and TH-GFP-ASYN-KO positive neurons (i.e. dopaminergic neurons) treated with 647-tagged ASYN or with vehicle were selected for analysis. To analyze the effect of ASYN-oligomers on mitochondrial membrane potential, the mean fluorescence intensity (MFI) of the TMRE signal was measured in the presence of ASYN-oligomers and normalized to the values obtained in the absence of ASYN-oligomers. To analyze the effect of ASYN-oligomers on the expression of TH-GFP, the MFI of TH was measured in neurons containing 647-tagged ASYN-oligomers vs. neurons without 647-tagged ASYN-oligomers. As the image analysis (FIJI) software provides automated MFI measurements, blinding was not essential.

### Immunocytology of neuronal cell cultures

4% PFA fixed neuronal cell cultures were washed 3×10 min in phosphate buffered saline (PBS), blocked using a blocking solution (BS) (0,2% Triton X-100 in PBS and 5% donkey serum (DS)) for 1 h at room temperature, and incubated with mouse anti-TH (1:500, Millipore) or rabbit anti-NeuN primary antibodies in BS overnight at 4 °C. On the next-day cells were washed 4×10 min with PBS, incubated with donkey Alexa® 555 anti-mouse or anti-rabbit secondary antibodies for 1 h at room temperature and washed 4×10 min with PBS.

### Quantification of neuronal death

The toxic effect of ASYN oligomers/monomers, paraquat and rotenone on dopaminergic neurons was assessed through manual counting of immunostained TH^+^ neurons after treatment. Dopaminergic TH+ neurons were observed using an inverted fluorescence microscope (Olympus) under a 20x objective. The diameter of every well was scanned in two perpendicular directions (i.e. top to bottom and left to right) and total TH+ neurons were counted for every well. In the case of NeuN^+^ cells the scan was performed on the perpendicular direction only (i.e. top to bottom).

### Seahorse experiments

A Seahorse Bioscience XF24 analyzer (Agilent Technologies, Santa Clara, CA, USA) was used to measure the mitochondrial O_2_ consumption rate (OCR) and the extracellular acidification rate (ECAR). To this purpose, 1×10^6^ primary midbrain neurons (E14.5) pooled from 3 pups were seeded per well on a Seahorse XF96 plate and cultured for 14 days in neuron growth medium (Neurobasal Medium, 2% B27, 0,25% Glutamax, 1% Penicillin/Streptomycin; all Thermo Fisher Scientific) with the different treatments before performing the Seahorse run.

The OCR and ECAR measurements were performed in Seahorse XF Assay medium (Agilent Technologies), which is an unbuffered medium allowing assessment of OCR and ECAR in parallel. 1 hr before the analysis, the neuron growth medium was removed, cells were washed once with XF Assay Medium and then supplied with 180μl XF Assay Medium freshly supplemented with 25 mM glucose. Subsequently, the cells were incubated at 37 °C under atmospheric CO_2_ and O_2_ concentrations. During the seahorse run the Oligomycin, Carbonyl cyanide-4 (trifluoromethoxy) phenylhydrazone (FCCP), Rotenone, Antimycin A were sequentially added reaching a final concentration of 1μg/ml, 1 μMm 2,5μM, and 2,5μM, respectively.

As the results of the seahorse measurements depend on the number of cells used, the results were normalized to cellular DNA content using the Quant-iT PicoGreen dsDNA Assay kit. Therefore, the plate to be normalized was frozen after the experiment. Frozen cell plates were thawed on ice and cells were subsequently lysed by adding 10 μl of Proteinase K (20 mg/ml) per well to the cells and their supernatant. After mixing, the plate was incubated for 1 h at 37°C and then for 10 min at RT. 10 μl of sample and standards were used for the assay with two replicates each. Next, the PicoGreen assay procedure was performed according to the manufacturer’s protocol. Fluorescence intensity was measured in microplates for fluorescence-based assays and DNA concentrations were calculated using linear regression analysis.

### Animal Fixation

At the end of all experiments (2 or 4 months timepoint), mice were deeply anesthetized using 200 mg/kg ketamine and 10 mg/kg xylazine and then transcardially perfused. After checking the posterior interventricular reflex and the eyelid closure reflex to ensure deep surgical anaesthesia, the thorax was opened and a button cannula was advanced into the left ventricle. The right atrium is opened to allow the blood to drain. After flushing the blood vessels for 5 min with PBS at a flow rate of 1 ml/min, perfusion fixation followed with 4% formaldehyde solution in phosphate buffer at a flow rate of 1 ml/min for 12 min.

### Statistical evaluation

For our *in vivo* and histology experiments, group size was predetermined with the help of our institute’s bio-statistician, using the following parameters: Significance level: α ≤ 0,05 / 6 = 0,00833 (Error 1. Type ≤ α), Power: 0,8 (Error 2. Type ≤ ß = 0,2), Effect size: f = 0,4. The program implemented for sample calculation and power analysis was: G*Power 3.1.3, Procedure “MANOVA” [23]. According to these calculations, our minimum group size was n=6. In order to be able to compensate for unforeseen problems (e.g. mortality), the group size was increased to n=7. *In vitro* experiments were performed in 96 well plates, and treatments were separated by column, so that all treatments of one experiment were performed in the same plate, under equal preparation and culture conditions. In order to avoid variability between preparations, we normalized to the control in each plate assuming that the conditions in which the neurons were prepared were homogeneous between all wells of the plate.

For the statistical analysis of our experiments we used GraphPad Prism 6.0, and a description of each statistical test used is provided in each corresponding figure legend. Graphs present individual data points wherever applicable. Datasets were assessed for normality (D’Agostino-Pearson normality test) and identification of outliers (ROUT method, Q = 1%; recommended by Graphpad). No outliers were identified. In all statistical tests, a p-value of < 0.05 was considered the threshold for declaring statistical significance.

## Results

### The absence of ASYN prevents the appearance of motor symptoms and the loss of dopaminergic neurons in the *substantia nigra* after oral exposure to rotenone

Our previous studies had shown that exposure to orally administered rotenone induces the appearance and progression of PD-like pathology from the ENS to the SN with loss of dopaminergic neurons following Braak′s staging in the course of 3 months [11, 12]. To investigate the role of ASYN in this progression, we exposed 1-yo ASYN KO mice (B6;129×1-*Snca*^*tm1Rosl*^/J, Jackson Laboratories) to orally administered rotenone or vehicle for 2 and 4 months (experimental timeline in Figure 1). WT (C57BL/6) mice where used as a control for the genetic background (results already published, see Figure 3 in [16]). In ASYN KO mice, the number of TH^+^ neurons was comparable to the number of dopaminergic neurons in the SNpc of WT mice and, contrary to what was observed in WT mice, orally administered rotenone did not induce changes in the number of TH^+^ neurons in the SN even after 4 months of rotenone exposure (see Figure 2A-F). Thus, suggesting that the presence of ASYN is necessary for PD pathology to progress. Interestingly, all ASYN KO mice groups showed a poorer performance in the rotarod test when compared to WT controls (Figure 2G). This would be in accordance with the alteration of the nigrostriatal dopamine system, despite a normal number of dopaminergic neurons in the SNc [24].

**Figure 1:**
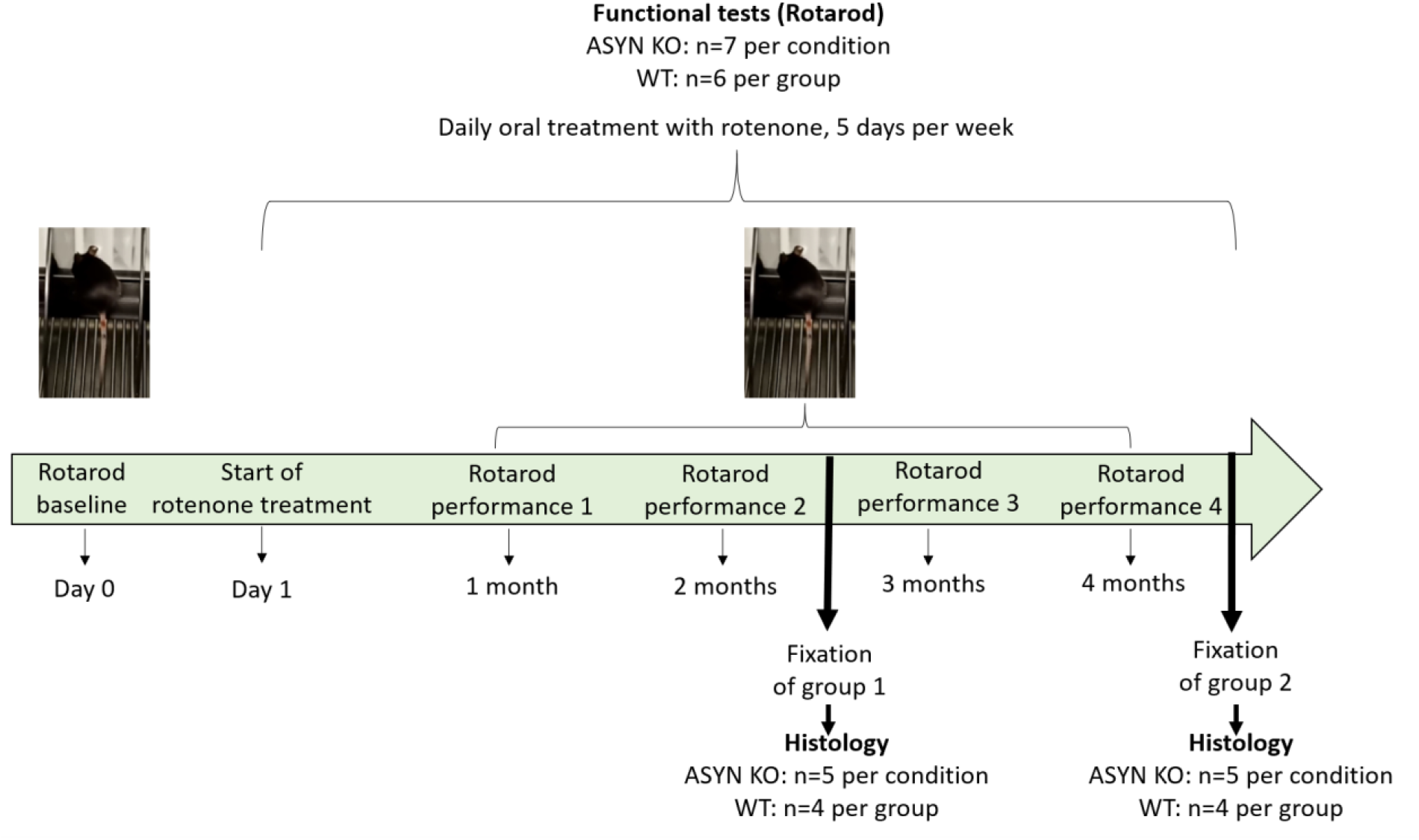
*In vivo* experimental timeline. Upon arrival and acclimatization, mice undergo the rotarod test for evaluation of motor performance before the start of rotenone treatment (Baseline, day 0). Following that, rotenone oral administration via administration by intragastric gavage was performed daily, 5 days per week for 2 (group 1) or 4 (group 2) months. Rotarod performance was assessed once every month until the end of the experiment (fixation).

**Figure 2:**
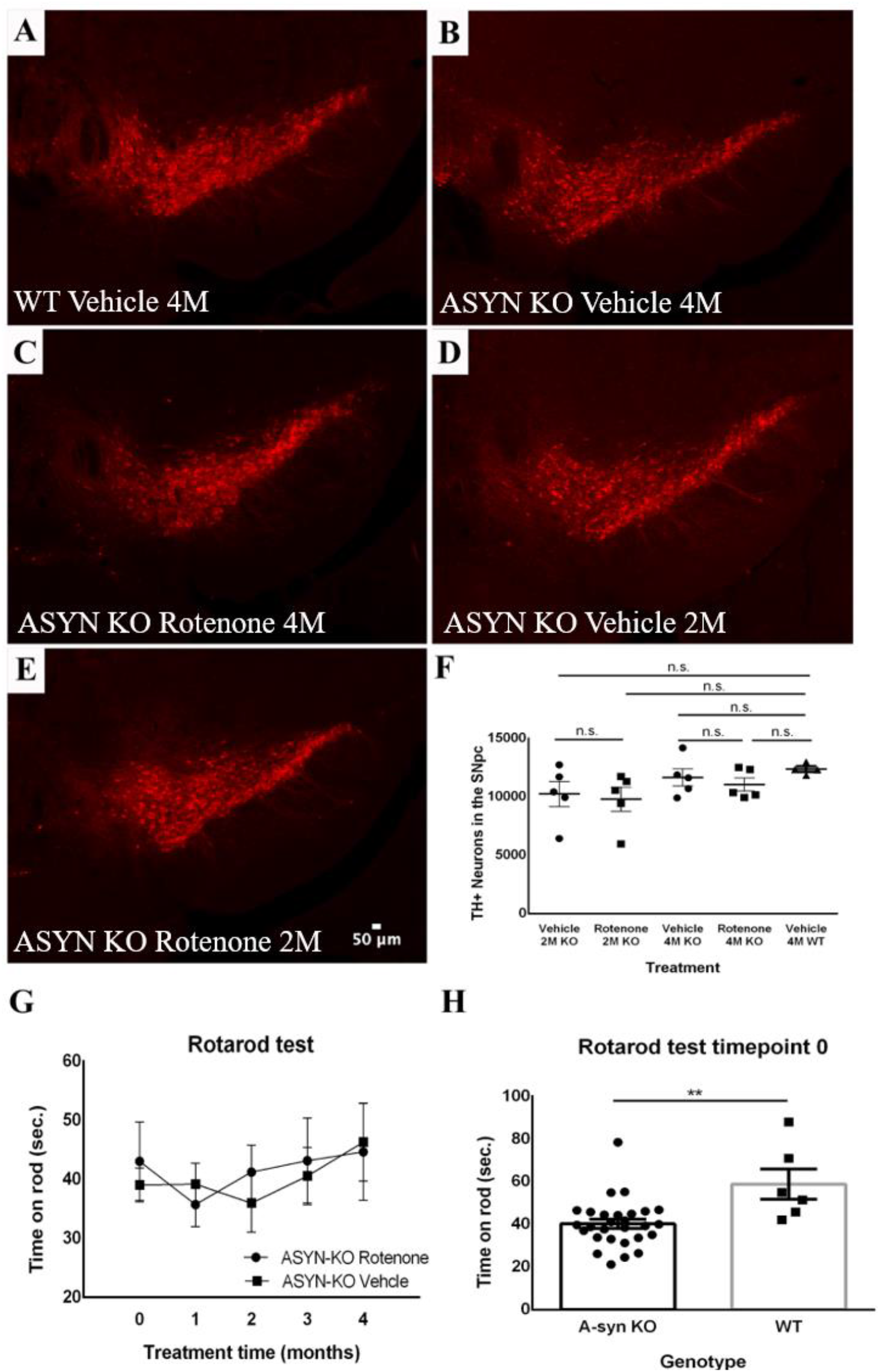
Effects of rotenone exposure on the number of Tyrosine Hydroxylase (TH^+^) neurons in the SNpc and motor function of alpha-synuclein knockout (ASYN-KO) and Wild-type (WT) mice. A-F: Representative confocal microscopy images showing TH+ neurons in the SNpc, labeled in red. A) WT Vehicle, B) ASYN KO Vehicle 4M, C) ASYN-KO Rotenone 4M, D) ASYN-KO Vehicle 2M, E) ASYN-KO Rotenone 2M. F) Graphic showing the average number of TH^+^ neurons in the SNpc in the different treatment groups of ASYN-KO mice. (1-way ANOVA: F=0.8964; p=0.4645, two-tailed unpaired Student’s ttests: p>0.05 for each comparison; ASYN KO: n=5, WT: n=4) G) Graphic showing the average rotarod performance across 4 months of Rotenone or Vehicle exposure. Treatment did not influence the motor performance of the mice at any timepoint (2-way ANOVA: F = 0.03337; P = 0.8581) H) Graphic showing the Rotarod performance of all ASYN KO mice vs WT mice at timepoint 0. WT mice were able to spend significantly more time on rod compared to ASYN-KO mice, independent of treatment (two-tailed unpaired Student’s ttest: mean diff.=18.57 ± 5.554; p=0.0021; ASYN KO: n=28, WT: n=6), *p<0.05; **p<0.01

**Figure 3:**
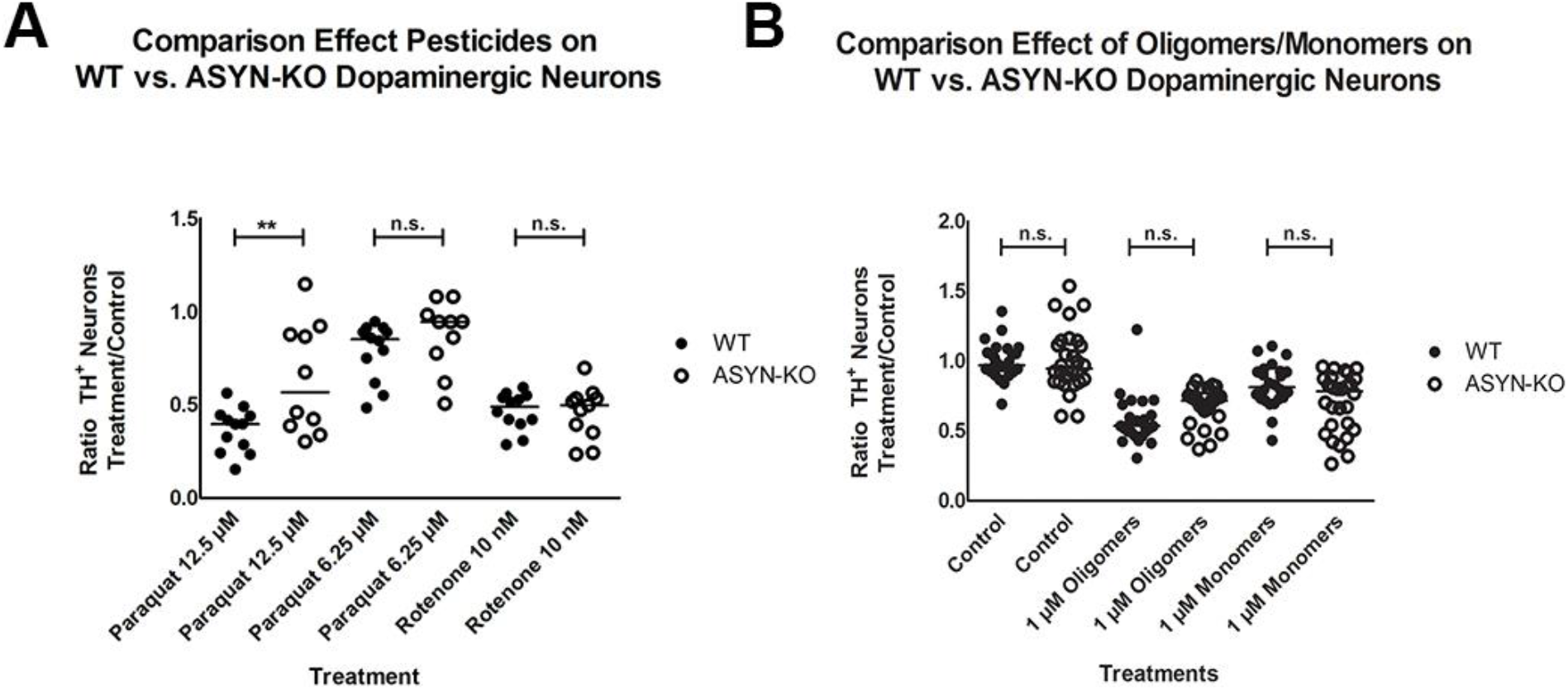
The presence of alpha-synuclein (ASYN) does not affect pesticide or ASYN monomer/oligomer dependent neuronal death. A) Significant increase in the survival of TH^+^ neurons in the ASYN-KO group compared to WT upon 12.5μM Paraquat exposure (two-tailed unpaired Student’s ttest; mean diff=0.274 ± 0.093, p=0.0081). No significant differences between ASYN KO and WT using lower Paraquat concentration (6.25μM) or 10nM Rotenone. B) No significant differences between the ASYNKO and WT cultures in the response to ASYN oligomer or monomer addition (two-tailed unpaired Student’s ttests: p>0.05 for each comparison). 6-8 embryos were used per experiment; each experiment was repeated at least 3 times. Each data point in the graphs represents the measurements from one well.

### The toxicity of rotenone and ASYN is independent of endogenous ASYN

It has been shown that the interaction between endocytosed ASYN oligomers with endogenously produced ASYN monomers induces the transformation of the latter ones into ASYN oligomers [25]. Different authors suggest that this prion-like mechanism amplifies the effect of the endocytosed ASYN and is responsible for its toxicity and the propagation of the PD pathology. In order to understand whether endogenous ASYN played a role in the toxicity exerted by paraquat, rotenone or ASYN on dopaminergic neurons, we compared the effect of paraquat (6.25 and 12.5 μM), 10 nM rotenone and 1 μM ASYN oligomers and monomers on primary mesencephalic neuronal cultures obtained from WT and ASYN KO mice. Our results show that the presence of endogenous ASYN did not influence the toxicity of 10 nM rotenone, 1 μM of ASYN oligomers or monomers and only showed a very moderate effect in the case of 12.5 μM of paraquat (Figure 3A and B).

### Interaction between endocytosed ASYN and host′s mitochondria is responsible for its toxicity

It has been shown that ASYN interacts with mitochondria. Therefore, in order to investigate whether this interaction could be responsible for ASYN toxicity, we analyzed mitochondrial activity in control and neurons treated with 647-tagged ASYN oligomers. For this, we labeled active mitochondria in mesencephalic neuronal cultures using tetramethylrhodamine ethyl ester (TMRE) in TH-GFP WT and TH-GFP/ASYN-KO cell cultures) after ASYN treatment. TMRE is a cell permeant, positively charged, red-orange dye that readily accumulates in active mitochondria due to their relative negative charge. Depolarized or inactive mitochondria have decreased membrane potential and fail to sequester TMRE. In order to differentiate TH^+^ neurons from the rest of neuronal subtypes during live cell imaging, we prepared mesencephalic primary cell cultures from TH-GFP mice and ASYN-KO/TH-GFP mice (Figure 4A-D). ASYN in TH-GFP^+^ neurons (arrowheads mark the soma of TH+ neurons in 3C and 3D) led to a drastic reduction of the mitochondrial membrane potential in those neuronal areas with 647-tagged ASYN inclusions (arrows in Figure 4C-D and quantification graph in 4E). We also observed that the intensity of the TH-GFP signal seemed to be reduced in neurons containing endocytosed ASYN oligomers. To test whether exogenous ASYN alters the expression of TH-GFP we measured the GFP signal fluorescence intensity in WT and ASYN KO TH-GFP^+^ neurons that showed no ASYN uptake vs WT and ASYN KO TH-GFP^+^ neurons that contained ASYN. Whereas TH^+^ neurons without ASYN oligomers showed no changes in the morphology or the intensity of the TH-GFP fluorescent signal (Figure 4A, B and F), TH^+^ neurons containing ASYN oligomers had a reduced TH-GFP fluorescent signal (* in 4C and D, quantification graph in 4F). Altogether, our results suggest that the toxicity of ASYN on mitochondria is proportional to the amount of endocytosed protein and to the proportion of mitochondria impaired by ASYN inclusions. Interestingly, whereas ASYN impaired mitochondria in all kind of cells in primary mesencephalic neuronal cultures, the number of NeuN^+^ neurons in mesencephalic cultures was not altered by exposure to ASYN (Supplementary Figure 2). Thus, suggesting that dopaminergic neurons were more susceptible to ASYN then other mesencephalic neurons.

**Figure 4:**
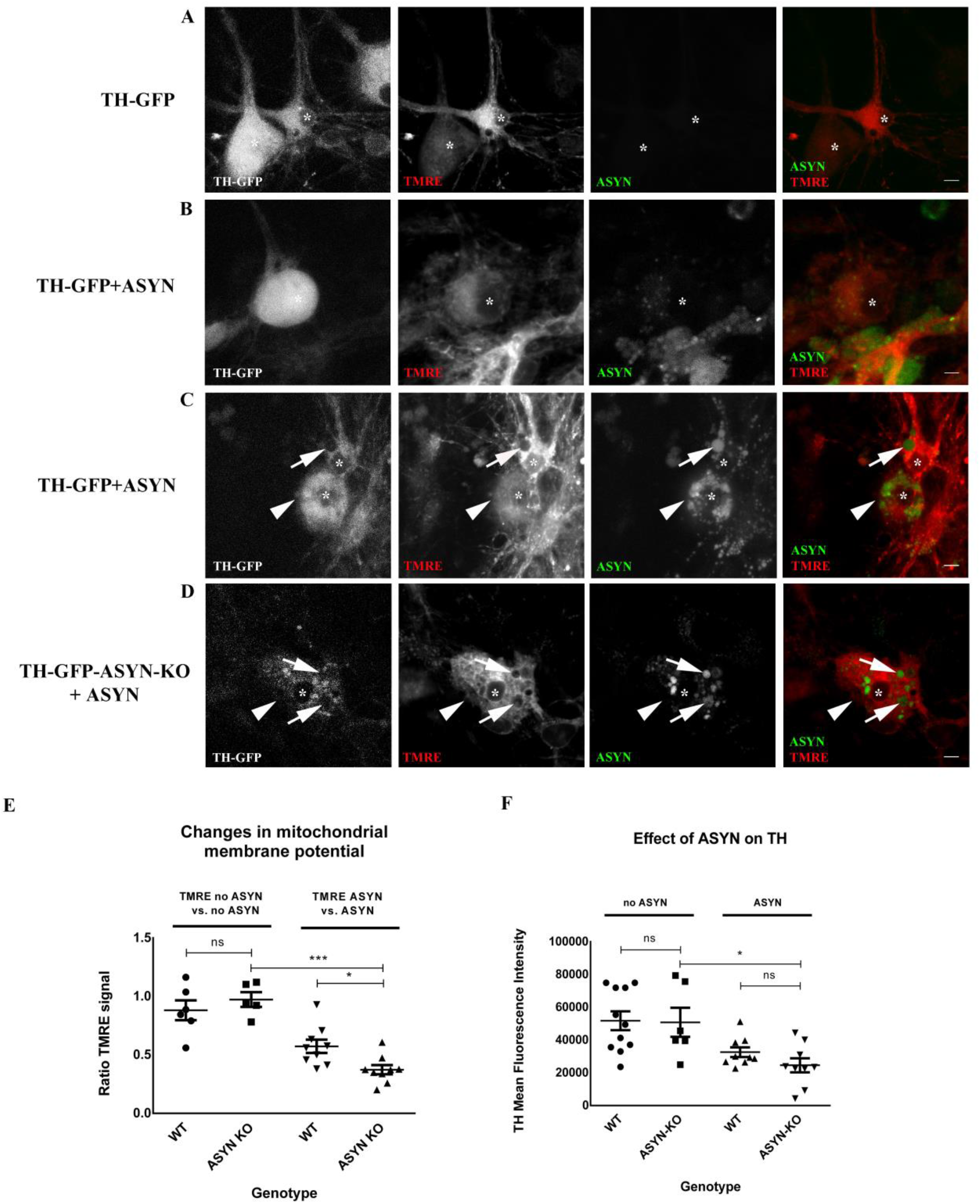
Alpha-synuclein (ASYN) oligomers induce the disruption of the mitochondrial network and impair mitochondrial membrane potential in dopaminergic neurons independent of endogenous ASYN: A, B, C and D show representative confocal images of TH-GFP WT and TH-GFP ASYN-KO neurons (*) treated with 647-ASYN oligomers (green) and stained with the live-dye TMRE (red). A) TH-GFP WT untreated neurons. B) TH-GFP WT neurons with low load of ASYN. C) TH-GFP WT neurons with higher amount of ASYN forming inclusions with no mitochondrial activity (arrows) and D) TH-GFP ASYN-KO neurons with high amounts of ASYN forming inclusions with no mitochondrial activity (arrows). Arrowheads in C and D mark TH-GFP expressing neurons. Scale bar: 20 μm. Dot-plot graphic in E shows the effect of 647 tagged ASYN-oligomers on mitochondrial membrane potential (TMRE signal). No significant difference in the TMRE signal ratio is observed between WT and ASYN KO for neurons without 647 tagged ASYN uptake (TMRE no ASYN vs. no ASYN) (1-way ANOVA followed by Newman-Keuls Multiple Comparison Test: mean diff.= 0.09206 ± 0.1099; p=0.4239). On the other hand, the 647 tagged ASYN -containing WT group has a higher TMRE signal ratio compared to 647 tagged ASYN -containing ASYN KO (1-way ANOVA followed by Newman-Keuls Multiple Comparison Test: mean diff.= -0.1999 ± 0.06873; p=0.0103). Finally, the ASYN KO neurons with 647 tagged ASYN inside have a significantly decreased TMRE signal ratio compared to the WT and ASYN KO neurons wihout 647 tagged ASYN inside (1-way ANOVA followed by Newman-Keuls Multiple Comparison Test: mean diff. = 0.6002 ± 0.06999; p= 0.0010) Dot-plot graphic in F shows the effect of 647-tagged ASYN oligomers on TH-GFP fluorescence intensity. No significant difference in the TH (GFP) mean fluorescence intensity is observed between WT and ASYN KO without ASYN uptake (no ASYN)(1-way ANOVA followed by Bonferroni’s Multiple Comparison Test: mean diff.= 949.5 ± 10159; p=0.9268). Similarly, no significant difference is observed between ASYN-containing WT and ASYN-containing ASYN KO (1-way ANOVA followed by Bonferroni’s Multiple Comparison Test: mean diff.= 7976 ± 5225; p=0.1464). However, there is a significant reduction in the TH signal in ASYN-containing ASYN KO and WT neurons compared to neurons without 647 tagged ASYN inside (1-way ANOVA followed by Bonferroni’s Multiple Comparison Test: mean diff.= 26221 ± 8900; p=0.0114). Each data point in the graphs represents the signal from 1 neuron.

### Distally uptaken ASYN is retrogradely transported and can impair mitochondrial membrane potential in the soma of sympathetic neurons

In order to test whether distally uptaken and retrogradely transported ASYN would have an effect on mitochondria located in the soma, we cultured sympathetic neurons using the Campenot chamber as previously described. These chambers allow to culture sympathetic soma in a different medium as their axons [11]. Using this ex vivo model, we administered labeled ASYN monomer and oligomers to sympathetic neurites and investigated their effect on the mitochondrial network (this time using mitotracker as a second compound to determine mitochondria membrane potential and avoid interference with Alexa-Fluor-488 tagged ASYN) of the soma in the other compartment. Our results show that up-taken and retrogradely transported ASYN oligomers but not ASYN monomers impair the membrane potential of the mitochondrial network (Figure 5).

**Figure 5:**
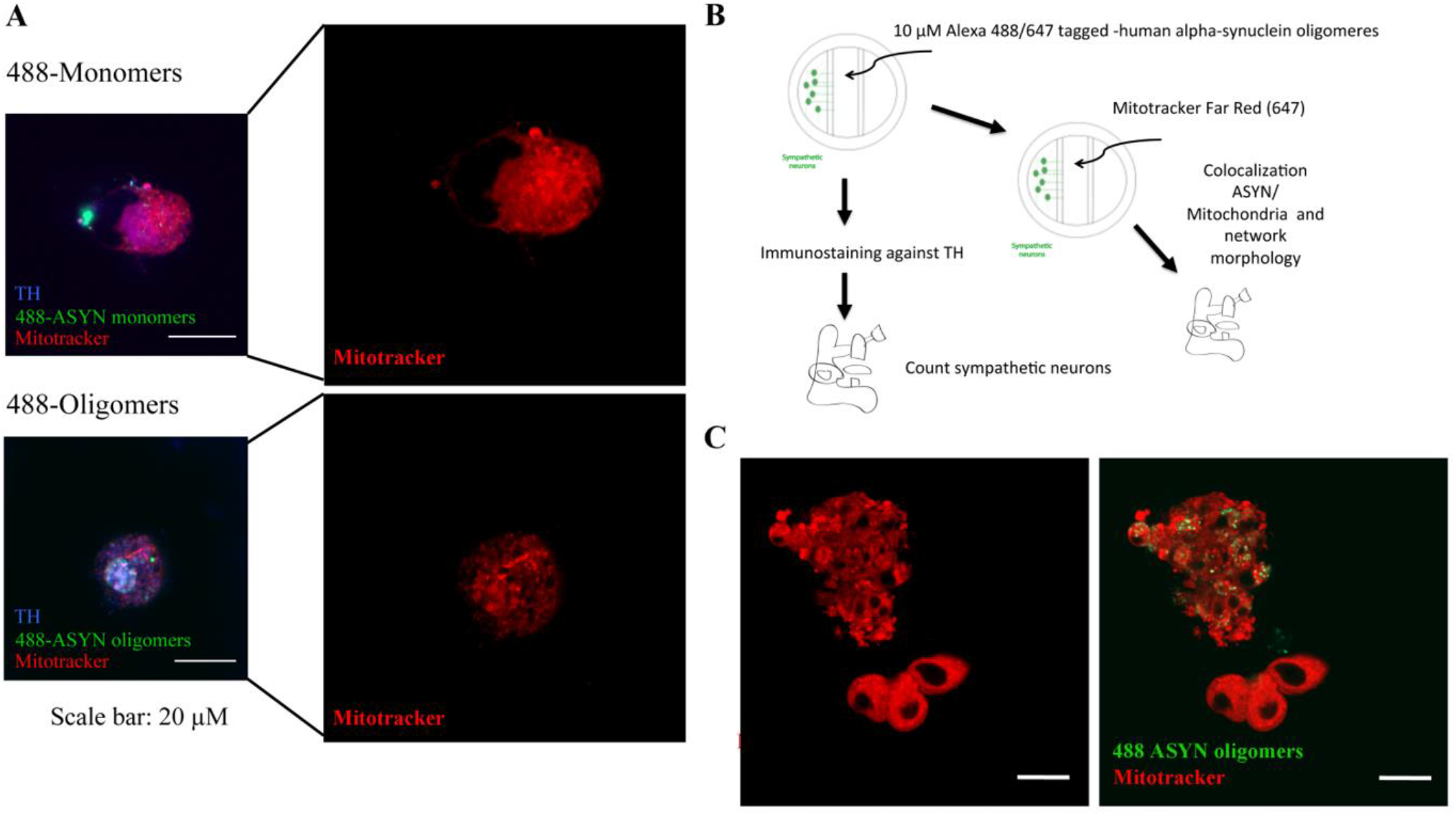
Effect of alpha-synuclein (ASYN) monomers/ oligomers on sympathetic neurons. A) Representative confocal images of sympathetic neurons treated with ASYN monomers (upper, green) or oligomers (lower, green) for one week. TH staining is shown in blue and the mitochodrial network is stained with Mitotracker (red) B) Schematic illustration of the experimental process C) Left: mitochondrial staining; Right: Colocalization of ASYN and mitochondrial staining. Scale bar in all images: 20 μm.

### ASYN KO mesencephalic neurons show a reduced basal respiration rate, a decreased ATP-linked respiration and a decreased maximal respiration

Determining the oxygen consumption rate (OCR) as well as the extracellular acidification rate (ECAR) of WT and KO neurons we did not detect any significant treatment effect neither when applying ASYN monomers or ASYN oligomers. However, there exists a clear-cut genotype effect both in respect to OCR as well as in respect to ECAR measurements. In detail, midbrain neurons derived from ASYN KO-mice show a decreased basal respiration rate (F(1,20) = 89,69; p<0,0001), decreased ATP-linked respiration (F(1,20) = 84,46; p<0,0001), decreased proton leak-linked respiration (F(1,20) = 149,1; p<0,0001), and decreased maximal respiration (F(1,20) = 75,73; p<0,0001). Also the spare respiratory capacity was reduced (F(1,20) = 75,73; p<0,0001). Furthermore, derived from the ECAR measurements it is evident that also basal glycolysis (F(1,20) = 161,0; p<0,0001), glycolytic capacity (F(1,20) = 89,69; p<0,0001) and the glycolytic reserve (F(1,20) = 230,4; p<0,0001) is significantly decreased in the KO neurons (Figure 6).

**Figure 6:**
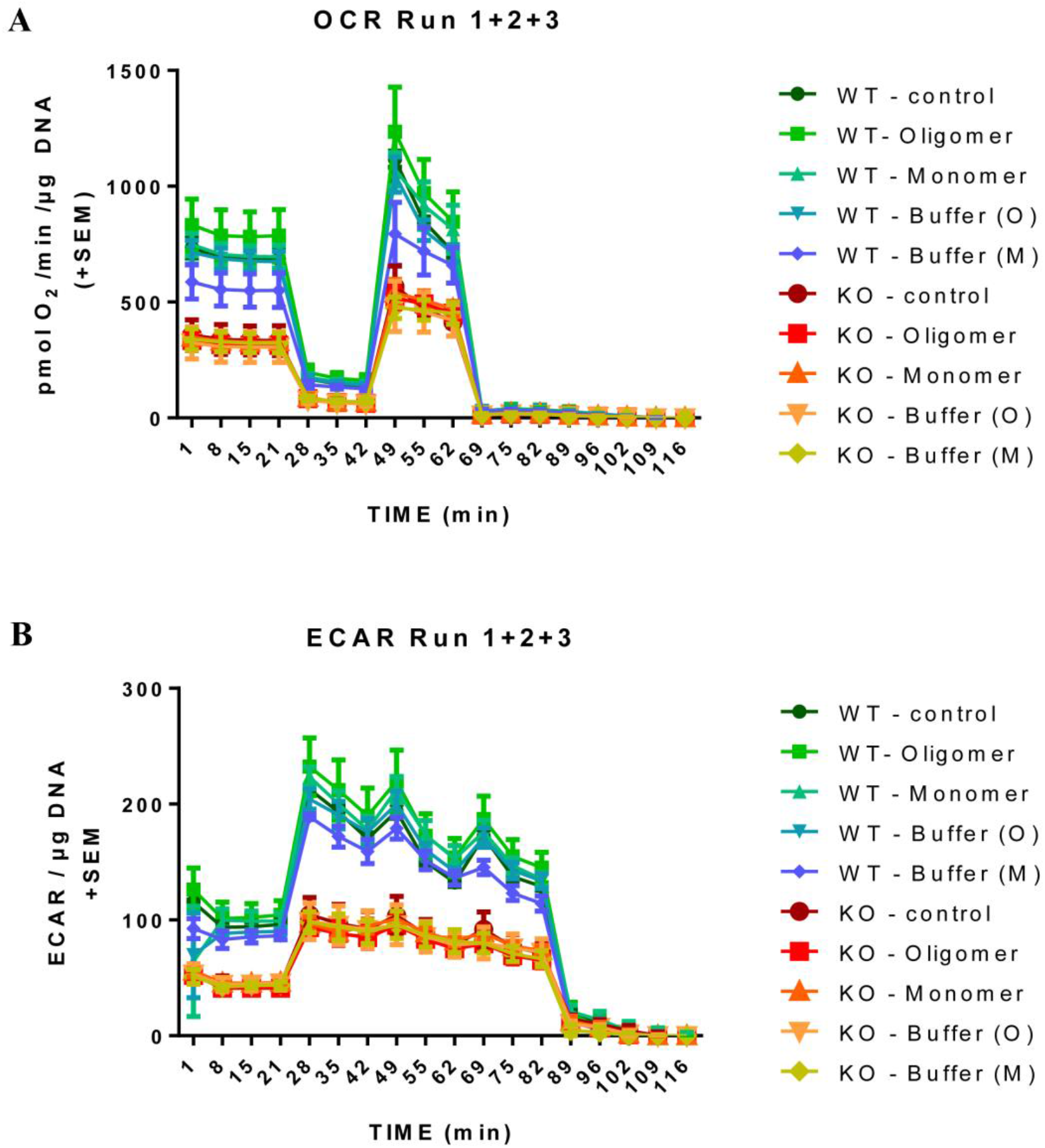
Effect of endogenous and exogenous alpha-synuclein (ASYN) on the Oxygen consumption rate (OCR) and extracellular acidification rate (ECAR) of mesencephalic neurons. OCR and ECAR measurements of WT and a-synuclein KO midbrain neurons with glucose as energy substrate challenged with mono- or oligomers of a-synuclein. OCR and ECAR analysis progresses from basal respiration, ATP- and proton leak-linked respiration after injection of oligomycin to maximal respiration after Carbonyl cyanide-4 (trifluoromethoxy) phenylhydrazone (FCCP) injection. Non-mitochondrial respiration resulting from RO injection has been subtracted from all values. (A) Analysis showed a genotype dependent decreased oxygen consumption rate throughout the experiment indicated by reduced OCR levels as well as (B) a genotype dependent decreased glycolytic activity. Treatments with mono- and oligomers did not have significant effects. Data were normalized to DNA content and n = 3 and 3 technical replicates were measured on each plate. Statistical analysis was done with n Prism 6.0 using a two-way Anova test.

### Dopaminergic neurons are more sensitive to rotenone and ASYN oligomers than cortical or enteric neurons

Finally, we asked whether the differences in the prevalence of non-motor vs. motor symptoms in patients could be explained by differences in the sensitivity to external *noxae*. For this we compared the sensitivity of different neuronal subpopulations that show ASYN pathology in PD patients during the progression of the disease to rotenone, paraquat and ASYN (oligomers and monomers). For this we exposed primary cortical, mesencephalic and enteric neuronal cultures to several concentrations of paraquat (6.25, 12.5 and 25 μM), 5 nM of rotenone or 0.1 and 1 of ASYN oligomers-using 0.1 and 1 μM of ASYN monomers as control. Surviving neurons were calculated as the number of NeuN^+^ neurons in the case of cortical neurons, as the number of TH^+^ neurons in the case of dopaminergic neurons and as the number of PGP9.5^+^ neurons in the case of enteric neurons. All results were normalized to the number of neurons in the control group. Cortical neurons were less sensitive than enteric or dopaminergic neurons (that had a similar sensitivity) to paraquat and dopaminergic neurons were much more sensitive to rotenone when compared to cortical or enteric neurons (Figure 7A-C). Interestingly, this was also the case upon exposure to ASYN. The highest concentrations of ASYN oligomers did not have any significant effect on cortical or enteric neurons -the latter showing a high variability of the results-but exposure to ASYN had a significant deleterious effect on dopaminergic neurons (Figure 7D-F). Interestingly, whereas 1 μM ASYN monomers did not have a negative effect on other neurons, they were able to induced cell death in dopaminergic neurons.

**Figure 7:**
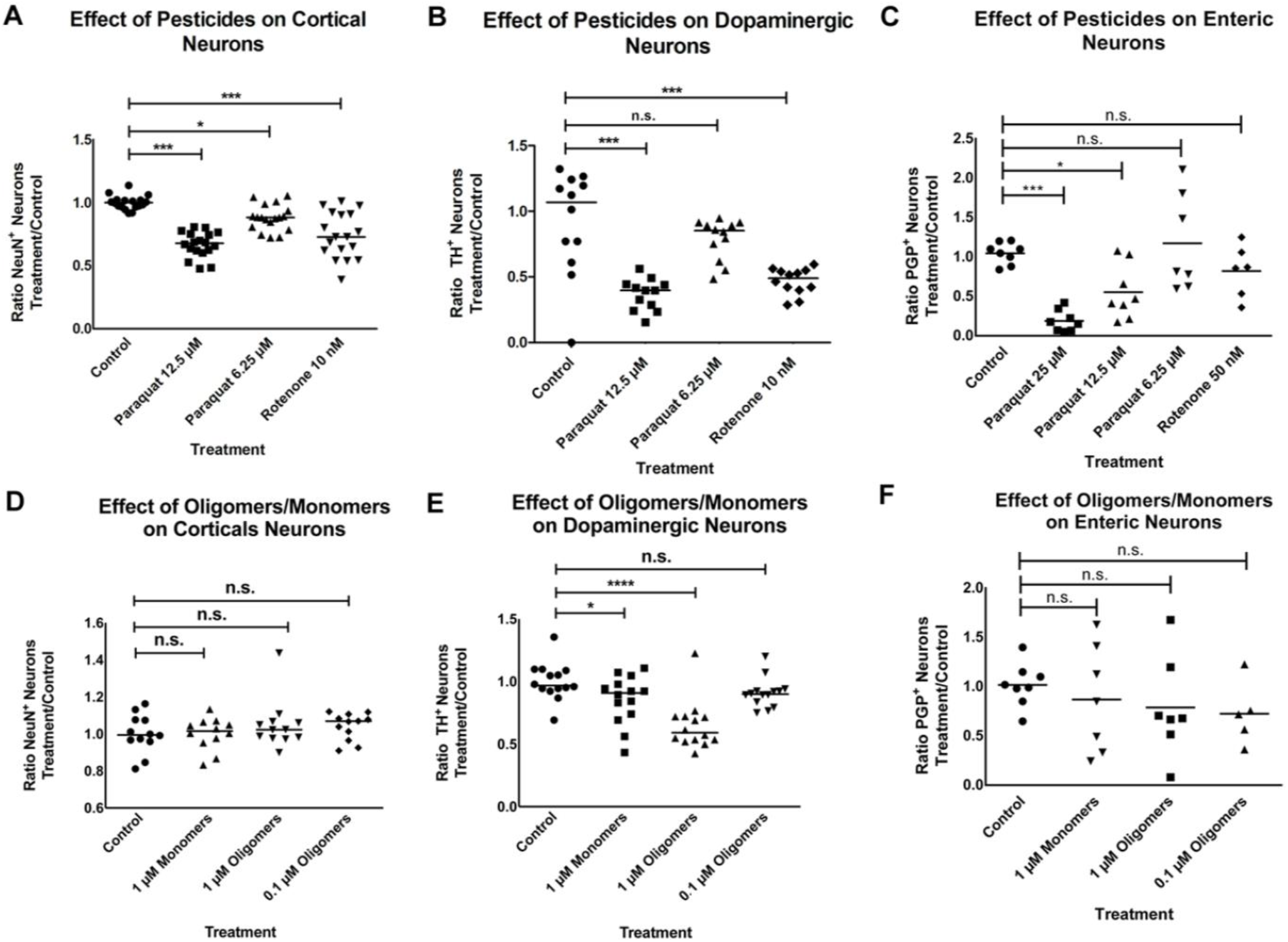
Effect of paraquat, rotenone or alpha-synuclein (ASYN) monomers/oligomers on different neuronal types. Cortical (A), Dopaminergic (B) and Enteric (C) neurons untreated (Control) or treated with Paraquat (12.5μM or 6.25μM) or Rotenone (10nM). The numbers of NeuN+ neurons counted for each treatment were normalized to the Control. A) In Cortical neurons, pesticide exposure had a significant effect on neuronal survival (1-way ANOVA: F= 27.93; p<0.0001). Dunnett’s multiple comparison’s test revealed significant neuronal loss compared to Control in the Paraquat 12.5μM and Rotenone 10nM treated cultures (p< 0,0001) as well as in the Paraquat 6.25μM treated cultures (p<0.01). B) In Dopaminergic neurons, pesticide exposure lead to a significant reduction of neuronal survival (1-way ANOVA: F= 15.86; p<0.0001). Dunnett’s multiple comparison’s test revealed significant neuronal loss compared to control in the Paraquat 12.5μM and Rotenone 10nM treated cultures (p< 0,0001) but not in the Paraquat 6.25μM ones (p>0.05). C) In Enteric neurons, pesticide exposure significantly decreased neuronal survival (1-way ANOVA: F=9.866; p<0.0001). Dunnett’s multiple comparison’s test revealed significant neuronal loss in the Paraquat 25μM (p<0.001) and 12.5μM (p<0.05) but not Rotenone 50nM (p>0.05) treated cultures compared to control cultures. Cortical (D), Dopaminergic (E) and Enteric (F) neurons untreated (Control) or treated with 1μM or 0.1μM ASYN oligomers, or 1μM ASYN Monomers. D) In Cortical neurons ASYN monomer or oligomer exposure had no significant effect on neuronal survival (1-way ANOVA: F=0.770; p=0.516). E) In Dopaminergic neurons ASYN monomer or oligomer exposure had a significant effect on neuronal survival (1-way ANOVA: F=32.45; p<0.0001). Dunnett’s multiple comparison’s test revealed significant neuronal loss compared to Control in the 1μM Monomers group (p<0.001), as well as in the 0.1μM (p<0.001) and 1μM (p<0.0001) Oligomers group. F) In Enteric neurons, exposure to monomers or oligomers did not have a significant effect on neuronal survival (1-way ANOVA: F= 0.6095; p= 0.6156). 6-8 embryos were used per experiment; each experiment was repeated at least 3 times. Each point in the graphs represents one well.

## Discussion

In this study we tested i) the role of endogenous ASYN in the progression of PD-pathology and the appearance of motor symptoms in vivo using ASYN KO mice exposed to orally administered rotenone, ii) the role of endogenous ASYN in the toxicity of extracellular ASYN and PD-associated pesticides, iii) the interaction of extracellular administered ASYN with the host′s mitochondria, iv) the susceptibility of different neuronal subtypes affected in the course of PD to rotenone, paraquat and ASYN and v) the effect of endogenous and extracellular ASYN on mitochondrial function. Our results suggest that, whereas the presence of endogenous ASYN is necessary for the progression of PD-like pathology in mice, the toxicity of extracellular ASYN-oligomers is independent of endogenous ASYN and is related to the interaction between ASYN and mitochondria. Furthermore, our results suggest that the lack of endogenous ASYN has an impact on the motor performance in vivo and on the mitochondrial respiratory function at the cellular level. Finally, our results suggest that the basis for the differences in the prevalence between PD-related non-motor symptoms and motor symptoms could be the difference in the susceptibility to external *noxae* (especially those affecting mitochondrial dysfunction and oxidative stress) between neuronal subtypes.

The results obtained by exposing ASYN KO mice to orally administered rotenone confirm that endogenous ASYN is necessary for the pathology to progress within the central nervous system (CNS). In this model, exposure to rotenone reduces the performance of the rotarod test. Histologically, the main pathological findings are ASYN phosphorylation and aggregation in several nervous structures together with the loss of dopaminergic neurons in the *substantia nigra* starting after 3 months of exposure [11, 12]. Interestingly, the progression of the pathology occurs between synaptically connected nervous structures. In ASYN KO we only could analyze the performance of the rotarod test and the loss of dopaminergic neurons in the *substantia nigra*, as there was no ASYN to be stained. Indeed, in the absence of ASYN, we did not observe any alteration in the performance of rotenone-exposed mice on the rotarod test or any loss of dopaminergic neurons even after 4 months of exposure to rotenone. Interestingly, when compared to WT mice, we observed that ASYN KO mice showed a poorer performance in the rotarod test even before exposure to rotenone. In WT mice, exposure to rotenone normally leads to impair motor function and a significant decrease in the number of dopaminergic TH^+^ neurons in rotenone-exposed mice when compared to vehicle-treated mice [11, 12, 16]. Whereas there seems to be a disconnection between the normal number of TH^+^ neurons in the SNc and the reduced performance in the rotarod test, these results would be in accordance with previous observations. An alteration of the nigrostriatal dopamine system with a normal number of dopaminergic neurons in the SNc has already been described in the literature [24]. As the authors write in their article “Here, we show that α-Syn−/− mice are viable and fertile, exhibit intact brain architecture, and possess a normal complement of dopaminergic cell bodies, fibers, and synapses. Nigrostriatal terminals of α-Syn−/− mice display a standard pattern of dopamine (DA) discharge and reuptake in response to simple electrical stimulation. However, they exhibit an increased release with paired stimuli that can be mimicked by elevated Ca^2+^. Concurrent with the altered DA release, α-Syn−/− mice display a reduction in striatal DA and an attenuation of DA-dependent locomotor response to amphetamine. These findings support the hypothesis that α-Syn is an essential presynaptic, activity-dependent negative regulator of DA neurotransmission.”

Remarkably, our data in several *ex vivo* cell culture models challenge the current accepted hypothesis that PD is a prion-like disease in which endocytosed ASYN oligomers serve as seeds for the aggregation of the host′s endogenous ASYN and are the main drivers of pathology progression [25-28]. If the seeding effect of endocytosed ASYN oligomers on the endogenous ASYN were crucial to the pathophysiological process and the appearance of PD pathology in host neurons, one would expect the lack of endogenous ASYN to reduce the toxicity of ASYN oligomers. Whereas we observed differences in the sensitivity to rotenone, paraquat and ASYN between cortical, enteric and dopaminergic neurons, we did not observe any differences in the sensitivity to these compounds between WT and ASYN KO neurons. Rather, the toxic effect of ASYN seems to be linked to a direct interaction with the host′s mitochondria. This would be in accordance with previously reported results in patients showing impairment of the mitochondrial respiratory chain and in several *ex vivo* models showing that ASYN can interact with mitochondria thereby impairing mitochondrial function. It has been shown that ASYN interacts directly with mitochondrial respiration complex I [29], can form pores in the mitochondrial membranes [30], increase the mitochondrial calcium concentration (reviewed in [31]) and can impair intracellular transport through the alteration of microtubules (reviewed in [32]). All these alterations can lead to alterations in the mitochondrial membrane potential, reduce energy production and increase oxidative stress, thereby increasing protein acetylation and oxidation. All this is especially important in PD, as it has been shown that there is a direct link between mitochondrial impairment, oxidation of iron and other metals, ASYN oxidation and phosphorylation and the aggregation of ASYN [33]. It has also been shown that ASYN can escape the cellular digestion pathway by rupturing lysosomes[34], which would explain their free interaction with mitochondria and the results obtained with sympathetic neurons when adding ASYN oligomers only to the sympathetic neurites using the Campenot chamber.

In terms of species to species variation and its effects on seeding activity, the literature is relatively scarce as most of the studies on seeding activity in mice are based on human ASYN overexpressing transgenic lines. In studies in which the question of different species can be assessed, the observations reported are relatively variable, due to different objectives as well as different experimental setups and conditions. While Rey et al. observed rapid internalization of human non-fibrillar ASYN into the olfactory bulb of WT-mice and spreading into interconnected brain regions without primary involvement of endogenous ASYN of the host (which was not the question addressed) [35], Barth et al. describe seeding effects of human ASYN fibrils in hippocampal slice cultures of WT mice as early as one week post-seeding indicating at least some interaction and seeding of ASYN between different species [36]. Although according to Fares et al. murine ASYN attenuates seeding and spreading induced by human ASYN PFFs [37]. Also Winner et al. investigated the effect of human ASYN in rats, showing that ASYN oligomers are more toxic than fibrils [38]. Finally, interaction and seeding activity between species can be observed in suitable models and with adequate observation timeframes but the extent may be limited by mixed species. *In vitro*, such a seeding effect between human ASYN and mouse and rat ASYN was described by Desplats et al. [27]. In this study, they show how ASYN is transmitted between neurons leading to inclusion formation and interactions with the hosts ASYN.

Altogether, our results suggest that the impairment of mitochondrial function through the interaction of endocytosed aggregated ASYN with the host′s mitochondria is key to the whole pathophysiological process driving the progression of PD within the CNS. Therefore, suggesting that the previously described seeding effect of ASYN plays, if at all, a marginal role in this process. All these findings and that of others suggest a different pathophysiological mechanism in PD progression. It has also been shown that extracellular ASYN is able to trigger an inflammatory reaction that contributes to the pathological process [39-42]. On the other hand, it has also been shown that inflammatory processes in the intestine also increase ASYN expression and can lead to aggregation [43, 44]. Both findings suggest that the interplay between ASYN and inflammation, and vice versa, also plays an important role in the appearance and progression of the disease.

We hypothesize that chronic exposure to environmental toxins or other chronical conditions (e.g. chronic intestinal inflammation as in the case of ulcerative colitis) leading to increased oxidative stress and ASYN aggregation triggers the appearance and progression of PD pathology. Initially aggregated ASYN is released by the neuron, up taken up by synaptically connected neurons and transported to the soma, where it interacts with the host′s mitochondria. This causes dysfunction of mitochondria and an increase in oxidative stress that leads to the oxidation, phosphorylation and aggregation of endogenous ASYN, that is again released by the host neuron. Released ASYN re-starts the cycle when up taken by synaptically connected neurons and triggers an inflammatory reaction.

We tried to quantify the effect of ASYN oligomers on mitochondria by measuring the OCR and the ECAR in primary mesencephalic cultures with the help of the Sea Horse system. However, the experiments performed using Sea Horse did not show any effect of ASYN monomers or oligomers on the OCR and ECAR of mesencephalic neuronal cultures. The lack of effect of ASYN oligomers on the total amount of NeuN^+^ mesencephalic cells, as opposed to the quantification of TH^+^ neurons, suggests that this could be due to the scarce presence of dopaminergic neurons in mesencephalic neuronal cultures. It has been reported that TH+ neurons represent 2-4% of the whole cell population on the coverslip in primary mesencephalic neuronal cultures [45]. We tried using Fluorescence Activated Cell Sorting (FACS) to enhance the proportion of dopaminergic neurons in cell cultures. However, TH-GFP dopaminergic neurons did not survive this process. Interestingly, ASYN KO neurons show lower reduced basal respiration rate, a decreased ATP-linked respiration and a decreased maximal respiration. This could, together with the above-mentioned alteration of the nigrostriatal dopamine system, explain the difference between WT and ASYN KO mice observed in the rotarod performance.

Finally, whereas PD-pathology, including the presence of aggregated ASYN in the form of Lewy bodies and Lewy neurites, can be observed throughout the nervous system, only motor symptoms and the loss of dopaminergic neurons are a consistent feature in PD patients. The prevalence of non-motor symptoms is as a whole equally consistent, however there is a high variability and inconsistency regarding the type of non-motor symptoms that PD-patients present and the type of neurons responsible for each symptom. For example, the prevalence of constipation is of 46.5% (range: 27.5–71.7%) for which alterations in cholinergic and sympathetic neurons as well as enteric neurons has been described [13, 14, 46]. In the case of the REM sleep behaviour disorder (RBD) the prevalence is of 34.2% with (range: 29.6–38.7%) and current research suggests that neuronal degeneration of brainstem nuclei, including the pontine tegmental area and medulla, is the pathophysiological cornerstone of RBD [7, 47]. Our current results comparing the effect of rotenone, paraquat and ASYN between neuronal subtypes suggest that this variability in the prevalence of motor and non-motor symptoms can be explained by the differences in the sensitivity to ASYN and oxidative stress between the different neuronal subtypes, with dopaminergic neurons being the most sensitive against both oxidative stress (caused by rotenone and paraquat [48-52]) and ASYN oligomers, which would explain why lack of dopamine-dependent motor symptoms are always present in PD patients. The effect of these *noxae* on enteric neurons is difficult to interpret, as we observed a high variability between experiments. We have previously shown that ASYN can be up taken by both enteric neurons (PGP 9.5^+^) and smooth muscle cells (SMA^+^) present in the cell culture [11]. Therefore, we believe that this high variability could be due to differences in the amount of non-neuronal cells between cultures, the presence of which is difficult to control.

The results of this study challenge the hypothesis that ASYN seeding is the main molecular mechanism underlying pathology progression and the appearance of the pathology in the host neuron and propose an alternative, more complex, molecular mechanism underlying the progression of PD in patients.

## Acknowledgements

This work was funded by the Deutsche Forschungsgemeinschaft (DFG, German Research Foundation) under Germany’s Excellence Strategy within the framework of the Munich Cluster for Systems Neurology (EXC 2145 SyNergy – ID 390857198). We would like to thank Fang Zhang with her help with the animal experiments. The work from Amit Khairnar was financed by a grant from the People Programme (Marie Curie action) of the Seventh Framework Programme of the EU according to the REA Grant Agreement No. 291782.

## Author′s contribution list

FP-M designed the study. YD performed animal experiments and TS performed the histology and the quantification of dopaminergic neurons in the SN. YD performed the *ex vivo* experiments with cell cultures. PD, YD and AS performed the Sea Horse experiments. PD, FG and DV-W analysed the Sea Horse data. VR and FS generated ASYN oligomers and monomers. AC, YD, FG, DV-W and FP-M wrote the manuscript. All other authors contributed conceptually and critically reviewed the manuscript.

## List of Abbreviations

ASYN: alpha-synuclein
PD: Parkinson’s Disease
BS: Blocking solution
CNS: Central nervous system
DIV: Day *in vitro*
DMV: Motor nucleus of the vagus
DS: Donkey serum
ECAR: Extracellular acidification rate
ENS: Enteric nervous system
FACS: Fluorescence Activated Cell Sorting
FCCP: Carbonyl cyanide-4 (trifluoromethoxy) phenylhydrazone
GFP: Green-fluorescent protein
IML: Intermediolateral nucleus of the spinal cord
KO: Knock-out
LB: Lewy bodies
LN: Lewy neurites
MFI: Mean fluorescence intensity
NeuN: Neuronal nuclei
NGF: Nerve growth factor
OB: Olfactory bulb
OCR: Mitochondrial Oxygen consumption rate
PBS: Phosphate buffered saline
PFA: Paraformaldehyde
PGP9.5: Protein gene product 9.5
RBD: REM sleep behaviour disorder
REM: Rapid eye movement
RO: Rotenone
RT: Room temperature
SNc: Substantia nigra pars compacta
TH: Tyrosine hydroxylase
TMRE: Tetramethylrhodamine, ethyl ester
WT: Wild-type

