## Supplemental Material for "Toxicity of extracellular alpha-synuclein is independent of intracellular alpha-synuclein"

[1] *Department of Psychiatry, Ludwig-Maximilian University Hospital, Nußbaumstr. 7, 80366, Munich*
[2] *Department of Neurology, Ludwig-Maximilian University Hospital, Marchioninistr. 15, 81377 Munich*
[3] *Center for Neuropathology und Prion Research, Ludwig-Maximilian-Universität München, Feodor-Lynen Str. 23, 81377 Munich*
[4] *Munich Cluster for Systems Neurology, SyNergy*
[5] *Institute of Developmental Genetics, Helmholtz Zentrum Munich, Germany*
[6] *Univ. Lille, Inserm, CHU Lille, UMR-S 1172 - JPArc - Centre de Recherche Jean-Pierre AUBERT Neurosciences et Cancer, F-59000 Lille, France*
[7] *Applied Neuroscience Research Group, CEITEC - Central European Institute of Technology, Masaryk University, Brno, Czech Republic*
[8] *Deutsches Zentrum für Neurodegenerative Erkrankungen, Feodor-Lynen Str. 17, 81377 Munich*
[9] *Technische Universität München-Weihenstephan 85764 Neuherberg/Munich, Germany*
[*] *These authors contributed equally and are listed in alphabetical order*
[#] *National Institute of Pharmaceutical Education and Research (NIPER), Ahmedabad, Palaj, Gandhinagar-382355, Gujarat, India*

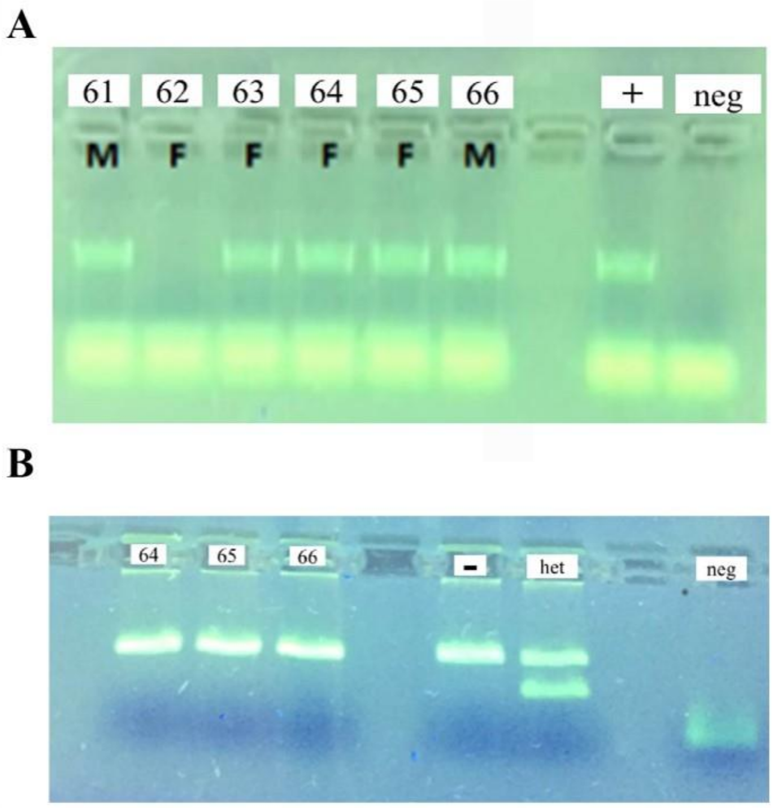


**Supplementary Figure 1: Alpha-synuclein knock-out (ASYN KO) and Green Fluorescent-Protein (GFP) genotyping**

Picture of genotyping results for GFP+ (A) and ASYN KO (B) mice. A) **+**= positive control, **neg** = negative control (water). B) **-** = ASYN KO mice: 192 bp (mutant), **het** = heterozygous mice: 105 bp (Wild-type) and 192 bp (mutant), neg = negative control (water). These results are in accordance to the expected results according to the provider.

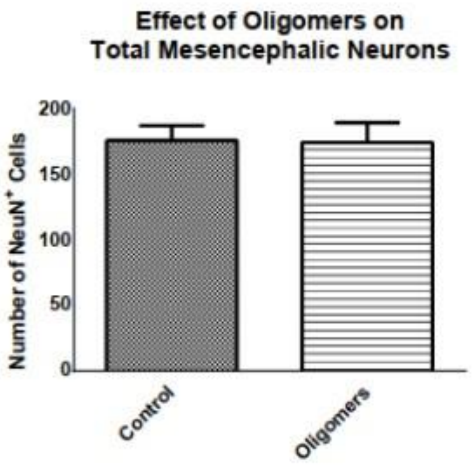


**Supplementary Figure 2: Effect of alpha-synuclein (ASYN) oligomers on the total amount of Neuronal nuclei-positive ($NeuN^+$) neurons in mesencephalic cultures.** Bar graph showing the total amount of $NeuN^+$ neurons in control treated and ASYN-oligomers treated mesencephalic cultures. The number in the y-axis shows the number of cells quantified in a vertical line from top to bottom for each well. Error bars represent standard error of mean (SEM).
